# 3D Passive Cavitation Mapping (3D-PCM) with a Large-aperture Planar Array

**DOI:** 10.64898/2026.06.20.733547

**Authors:** Chaorui Qiu, Daiwei Li, Haoming Huo, Arpit Mishra, Chenhang Li, Kaiping Yin, Nanchao Wang, Junqin Chen, Rui Yao, Ezra J Margolin, Michael Lipkin, Pei Zhong, Xiaoyue Ni, Junjie Yao

## Abstract

Urinary stone disease is a common urological condition with increasing incidence, particularly in developed countries. Laser lithotripsy (LL) has become a preferred minimally invasive treatment due to its high precision and low tissue damage. Recent studies suggest that cavitation plays a critical role in stone damage during LL, and three-dimensional passive cavitation mapping (3D-PCM) has emerged as a promising tool for detecting these events. However, clinical translation of 3D-PCM remains challenging due to limitations in imaging depth, field of view (FOV), and procedural compatibility. Here, we present a large-FOV dual-modality imaging system (3D-PCM and B-mode ultrasound) based on a large-aperture planar ultrasound array. Through array optimization and model-based reconstruction, our system achieves an expanded FOV of ∼40 × 40 mm^2^ at a clinically relevant imaging depth of ∼110 mm, while maintaining high spatial resolution of ∼0.6 mm laterally and ∼0.4 mm axially. *In vivo* experiments in a porcine model demonstrate that the reconstructed cavitation distribution correlates well with stone damage. Our technology has the potential to provide real-time treatment feedback during LL without disrupting the standard workflow.

## I. INTRODUCTION

Urinary stone disease is a common urological disorder of growing public-health concern. Stones forming in the ureter or kidney cause severe pain and trigger inflammatory responses that substantially affect patients’ quality of life. The disease affects ∼12% of the global population [1], [2], with even higher prevalence in developed countries, likely reflecting the contribution of metabolic syndrome [3]. In the United States, ∼1 in 11 individuals develops urinary stone disease in their lifetime, with prevalence nearly doubling over the past 15 years [4].

Laser lithotripsy (LL) has emerged as a mainstay minimally invasive therapy, offering greater targeting precision and less collateral injury than traditional shock wave lithotripsy [5]–[7]. During LL, an optical fiber is inserted into the urinary tract under ureteroscopic guidance to deliver short laser pulses (typically from a Ho:YAG laser) that progressively fragment the stone (Fig. 1a) [8]–[12]. In dusting mode (low energy, 0.2– 0.4 J; high repetition rate, 12–100 Hz), LL fragments the stone into fine debris, shortening procedure time and improving surgical efficiency [11], [13]–[15].

**Fig. 1.**
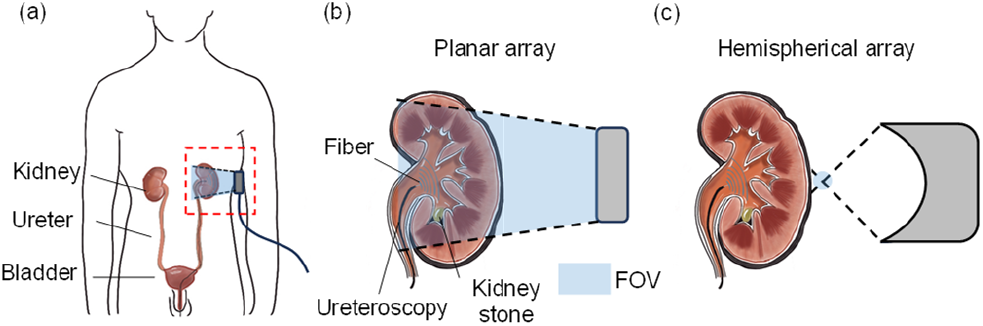
Schematic diagrams of (a) 3D-PCM during ureteroscopic laser lithotripsy. Comparison of the field of view (FOV) of (b) a planar array and (c) a hemispherical array.

**Fig. 2.**
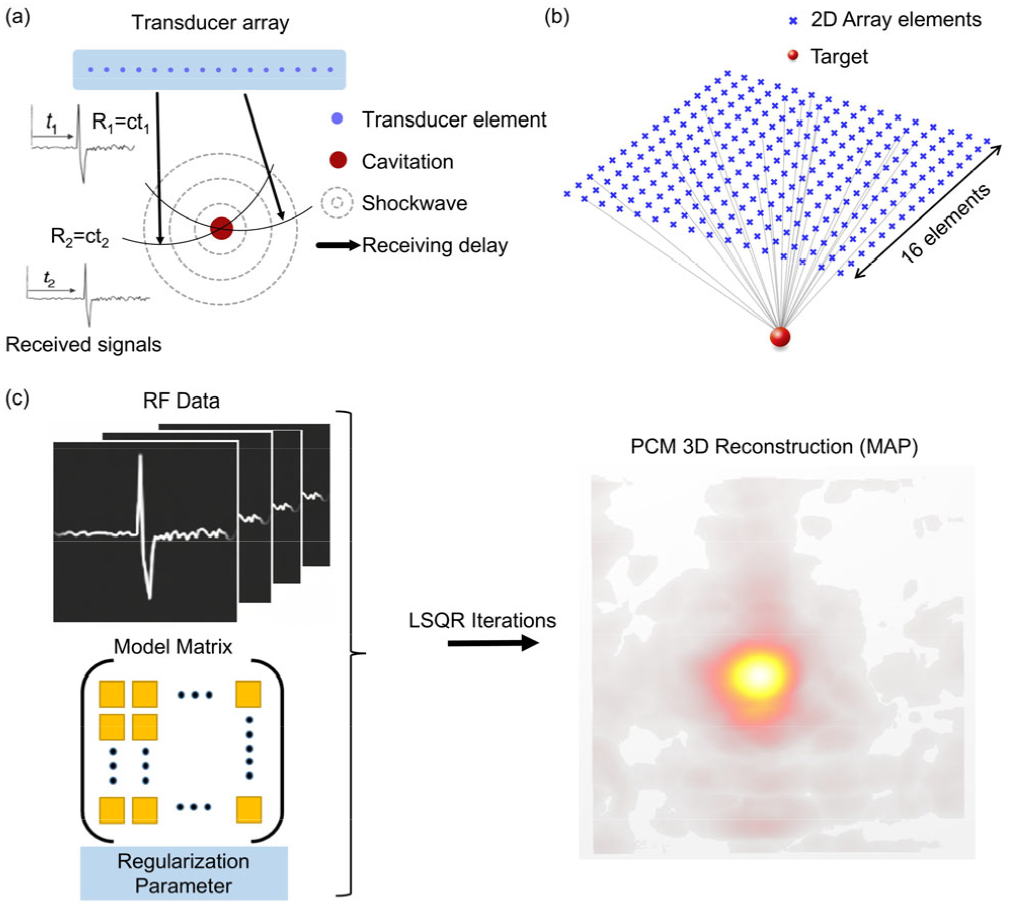
Model-based image reconstruction in 3D-PCM. (a) Geometry of the array elements relative to the target. (b) DAS beamforming for cavitation localization. (c) Model-based reconstruction pipeline and the corresponding 3D reconstruction image. The hot map cloud represents the intensity of the 3D reconstructed cavitation volume. LSQR, sparse linear equations and least square problems. MAP, maximum amplitude projection.

Although multiple pathways contribute to stone damage in dusting-mode LL, mounting evidence points to cavitation as a major and possibly dominant factor [13], [16], [17] [18]–[22]. Our own work revealed a strong correlation between stone damage and the cumulative spatiotemporal distribution of LL-induced bubbles, motivating real-time cavitation monitoring to optimize therapeutic outcomes [23]. However, current medical imaging approaches (e.g., B-mode ultrasound imaging) do not provide real-time three-dimensional (3D) visualization of cavitation at kidney-relevant depths [23]–[25]. A clinically viable solution must be non-ionizing, provide a field of view (FOV) large enough to localize bubble collapse events, and integrate seamlessly with standard LL workflows.

Recently, Li et al. developed a 3D passive cavitation mapping (3D-PCM) technique using a 256-element hemispherical ultrasound array. Owing to its high spatial resolution and receiving sensitivity, the system can resolve cavitation bubble collapse events with clear spatiotemporal detail. However, the hemispherical geometry of the array inherently restricts the field of view (∼8 × 8 × 8 mm^3^) and imaging depth (∼30-40 mm), posing a major barrier to clinical translation (Fig. 1c).

To address these limitations, we propose a system that enables 3D-PCM and B-mode imaging using a large-aperture planar ultrasound array (Fig. 1b). Through array optimization and model-based image reconstruction, our system substantially expands the FOV (∼40 × 40 mm^2^) and imaging depth (∼110 mm) while maintaining the spatial resolution (0.60 ± 0.06 mm lateral, 0.37 ± 0.07 mm axial) needed to resolve individual bubble collapse events, as confirmed by high-speed camera recordings. We further validated this approach in an *in vivo* porcine model, where the reconstructed cavitation distribution showed clear spatial correlation with stone damage. Collectively, these results suggest the clinical potential of the proposed technology to provide real-time procedural feedback during LL, improve surgical efficiency, and ultimately enhance patient outcomes.

## II. METHODS

### A. Image reconstruction of 3D-PCM

To improve reconstruction accuracy, we developed a model-based delay-and-sum (MB-DAS) reconstruction method. Model-based methods cast image reconstruction as an optimization problem in which the image is obtained by minimizing an objective function [26], [27]. Compared with delay-and-sum (DAS) beamforming, model-based methods offer greater flexibility and can mitigate several well-known limitations of analytical reconstruction [27]–[29]. The objective function typically combines a data-fidelity term with regularization terms encoding prior assumptions about the image, and the solution may be further constrained by a suitable basis [30], [31]. Specifically, model-based reconstruction minimizes the discrepancy between measured acoustic signals and those predicted by a forward model. Discretizing the linear forward model for acoustic wave propagation yields

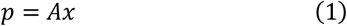

where *p* denotes the PCM signal, *x* denotes the initial pressure distribution within the tissue, and *A* is the model matrix, which depends only on the system geometry and the speed of sound in the tissue. The solution of Eq. (1) is obtained by minimizing the squared error:

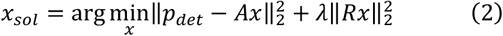

Here, *R* denotes the regularization operator, *λ* > 0 is the regularization weight, and ‖·‖_2_ represents the *L*^2^ norm. In practice, it is generally infeasible to explicitly compute the inverse of *A*. Instead, the solution is obtained through iterative numerical schemes; LSQR (least squares for sparse linear equations) is a common choice [32]. For limited-view detection geometries, the optimization problem in Eq. (2) is inherently ill-posed, making regularization essential for obtaining a stable solution [33], [34].

### B. Design of the planar ultrasound array

We performed acoustic field simulation with Field II to optimize the design of the 2D planar array [35], [36]. Under the linear time-invariant assumption, the emitted pressure field of the transducer at a field point ***r*** can be modeled as the convolution of the electrical excitation, the transducer electro-mechanical impulse response, and the spatial impulse response:

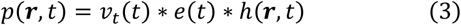

where *v*_t_(*t*) is the transducer electro-mechanical impulse response, *e*(*t*) is the applied voltage excitation, and *h*(***r***, *t*) is the spatial impulse response, which accounts for aperture geometry, focusing, and element distribution. A 1.5-cycle, 5-MHz square-wave voltage was used as excitation.

The transducer impulse response *v*_t_(*t*) was derived using the Krimholtz-Leedom-Matthaei (KLM) equivalent circuit model [37], which accounted for the element-size-dependent variations in sensitivity and bandwidth [38]. During simulation, the array’s transmission focus was steered to different depths and lateral positions. Based on the acoustic reciprocity principle, the receiving sensitivity pattern of the array is equivalent to its transmitting beam pattern under the same aperture and focusing conditions [36], [39]; we therefore assessed the array’s spatial resolution and detection sensitivity using the full width at half maximum (FWHM) and the peak intensity of the focused transmitting beam profile [40], [41].

### C. Fabrication and characterization of the planar array

Fig. 3 illustrates the fabrication process of the planar array. A PZT5H/epoxy 1-3 composite (Del Piezo Specialties LLC, USA) with a resonance frequency of 5 MHz was selected as the active material. The composite has a 50% volume fraction and an electromechanical coupling coefficient of 62%. Cr/Au was sputtered on the surfaces of the 1-3 composite to form electrodes. To improve bandwidth and sensitivity, we added quarter-wavelength dual matching layers. The acoustic impedances of the high- and low-impedance matching layers (*Z*_*h*_ and *Z*_*l*_) were calculated according to the following equation [42]:

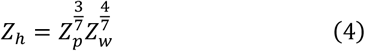

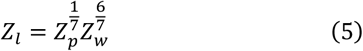

where *Z*_*p*_ and *Z*_*w*_ denote the acoustic impedance of the piezoelectric material and water, respectively. For the high-impedance matching layer, tungsten powder was mixed into epoxy (Epo-tek®301, Epoxy Technology, Inc., USA), tuning the acoustic impedance from 3.0 to 5.9 MRayl. For the low-impedance matching layer, polycarbonate (PC) with an impedance of 2.7 MRayl was used [43]. The matching layers were bonded sequentially using epoxy, and the thickness of the bonding layer was controlled to < 1 μm by applying pressure during curing. To electrically isolate and independently address each element, we used laser hatching technique (ProtoLaser R4, LPKF, Germany) to pattern the back electrodes on the 1-3 composite. A customized 256-channel double-layer flexible printed circuit board (FPCB; ∼80 µm thick) was then epoxy-bonded to the patterned electrodes on the 1-3 composite, where each solder pad was electrically and mechanically connected to a corresponding transducer element. Finally, a backing layer (thickness: ∼15 mm), composed of tungsten powder, hollow glass bubbles, and epoxy, was bonded to the rear side of the FPCB to suppress acoustic reflections and improve signal clarity.

**Fig. 3.**
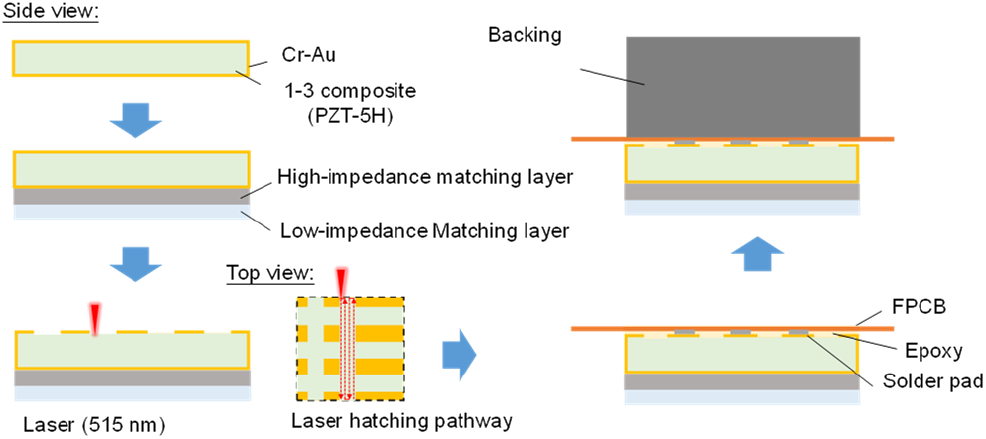
Fabrication process of the planar array.

The electrical impedance of the transducer element was measured using an impedance analyzer (4294A, Agilent Tech. Inc., USA). The center frequency and bandwidth were characterized by using an ultrasonic pulser/receiver (5900PR, Olympus Panametrics, USA). During the pulse-echo test, the transducer was excited with a 100 V electrical pulse, and the received echo signal was recorded without additional gain. Spatial resolution was characterized by imaging a wire-cross phantom composed of 100-µm-diameter tungsten wires using a conventional synthetic aperture method [44].

### D. Validation of 3D-PCM with high-speed camera

To validate our 3D-PCM system, we performed side-view shadowgraph imaging with a high-speed camera (Vision Research, Wayne, NJ) at 90,909 Hz to capture bubble collapse events induced by a single laser pulse from a Ho:YAG laser (Dornier MedTech, Munich, Germany). A 365-µm-core fiber was placed on top of a glass block (parallel to the glass surface, with a 1-mm standoff distance) to generate bubble dynamics. The LL laser pulse firing, high-speed camera recording, PCM signal acquisition, and 3D B-mode ultrasound imaging were precisely synchronized. B-mode ultrasound images were acquired at a 15-Hz frame rate for guidance, and a 3D ultrasound image was acquired prior to the LL treatment to localize the fiber tip and the glass surface. The planar array was placed 30 mm above the fiber tip.

A 10-Hz master trigger was generated and sent with each LL laser pulse to synchronize the high-speed camera and 3D-PCM acquisition. The 3D-PCM recording started 300 µs after the trigger and lasted for 800 µs to capture the full cavitation activity. For validation of the 3D-PCM results, the high-speed camera acquired 90 frames over a 1-ms window following the trigger, providing frame-by-frame visualization of the bubble dynamics.

### E. In vivo experiment protocol on a swine model

All animal procedures were approved by the Institutional Animal Care and Use Committe (IACUC; Protocol No. A071-24-03) [45], [46]. Five female Yorkshire pigs (75–100 lbs) underwent ureteroscopic LL under general anaesthesia. Pre-weighed cylindrical soft BegoStone phantoms (BEGO™, Lincoln, RI, USA) were fabricated with a 5:2 powder-to-water ratio and soaked in water for 24 h prior to implantation to achieve mechanical properties comparable to those of human urinary calculi [45], following our prior protocol [45].

Following cystoscopy and bilateral ureteral catheterization, a laparotomy and proximal ureterotomy were performed for stone implantation. Two stone phantoms were sequentially implanted into separate, non-adjacent renal calyces in each kidney. The ureterotomy was closed using 4-0 absorbable sutures, and watertight closure was confirmed. An 11/13 Fr ureteral access sheath was then introduced over a guidewire, and a single-use flexible ureteroscope (AXIS II, Dornier MedTech, Germany; OD 8.7 Fr; sheath-to-scope ratio 1.26) was advanced through the sheath to access the collecting system. To monitor local thermal response, K-type thermocouples (5SRTC-TT-K-36-36, OMEGA, Norwalk, CT, USA; 0.13-mm sensor diameter) were inserted through the renal parenchyma into the calyceal lumen under endoscopic guidance. The thermocouple tip was positioned adjacent to the stone-tissue interface, and the thermocouple-to-stone distance was maintained as consistently as possible across experiments. Endoscopic video recordings were acquired throughout the procedures.

LL was performed using a thulium fiber laser (TFL) (IPG Photonics, Marlborough, MA, USA) delivered through a 200-µm silica fiber (NA = 0.22). A dusting strategy was employed to reduce the stone to fragments smaller than 0.25 mm. LL continued to proceed until the stone was completely dusted.

Based on prior benchtop studies demonstrating favorable safety and fragmentation efficiency, laser treatments were performed at a pulse energy of 0.8 J and a repetition rate of 12 Hz (9.6 W) [46]. All procedures used room-temperature 0.9% saline irrigation continuously at a flow rate of 20 mL/min, controlled and delivered using a peristaltic pump (Masterflex®). TFL treatments consisted of alternating 30-second cycles of laser activation and 30-second rest until the stone was completely dusted, thereby limiting cumulative thermal loading.

Sham procedures were performed for 5 minutes in a predetermined alternating sequence and replicated all procedural steps, including ureteroscopy, irrigation, thermocouple placement, and temperature monitoring, without laser activation.

The entire procedure was repeated in the contralateral kidney. Upon completion of all experiments, animals were euthanized according to institutional guidelines. Kidneys were harvested and fixed for gross pathological and histopathological examination of laser-induced tissue injury.

#### In vivo 3D-PCM during laser lithotripsy

Fig. 4 illustrates the experimental setup for *in vivo* 3D-PCM during LL. The planar ultrasound array was connected to a programmable ultrasound scanner (Vantage 256, Verasonics; 19.235-MHz sampling rate), mounted on a robotic arm, and positioned over the kidney of an anesthetized swine (Fig. 4a). A flexible ureteroscope and laser lithotripter were introduced via the urethra to fragment the renal stone, while 3D-PCM monitored cavitation activity within the kidney in real time. Intraoperative photographs (Fig. 4b) show the robotic arm stabilizing the transducer array over the kidney, with ultrasound gel ensuring reliable coupling.

**Fig. 4.**
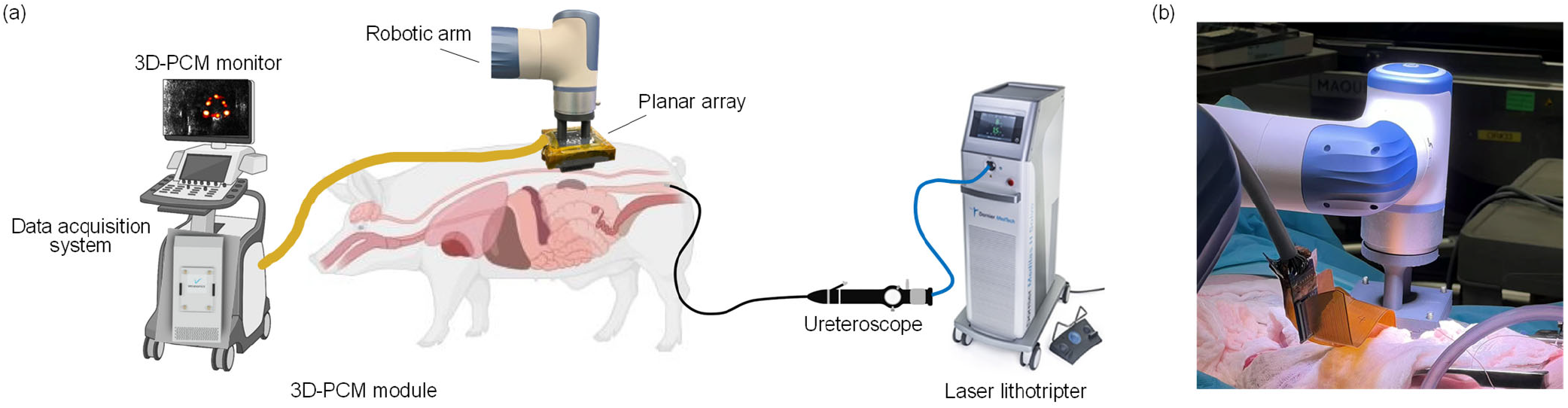
Experimental setup of in vivo 3D-PCM during LL treatment in a swine model. (a) Schematic diagram of the imaging setup. (b) Photo of the planar transducer array mounted on the robotic arm on top of the swine kidney.

## III. Results

### A. Simulation and design of the planar ultrasound array

To reconstruct the 3D cavitation distribution from 3D-PCM, the array must provide sufficient spatiotemporal resolution to localize individual bubble-collapse events; therefore, we used the FWHM of the beam profile to evaluate localization accuracy. Fig. 5b presents the FWHM as a function of imaging depth for 2D arrays operating at different frequencies, with element and pitch sizes equal to one wavelength. Although axial resolution improves with increasing frequency, lateral resolution shows no improvement and instead degrades with depth. This indicates that simple frequency scaling is not an effective strategy for improving localization performance.

**Fig. 5.**
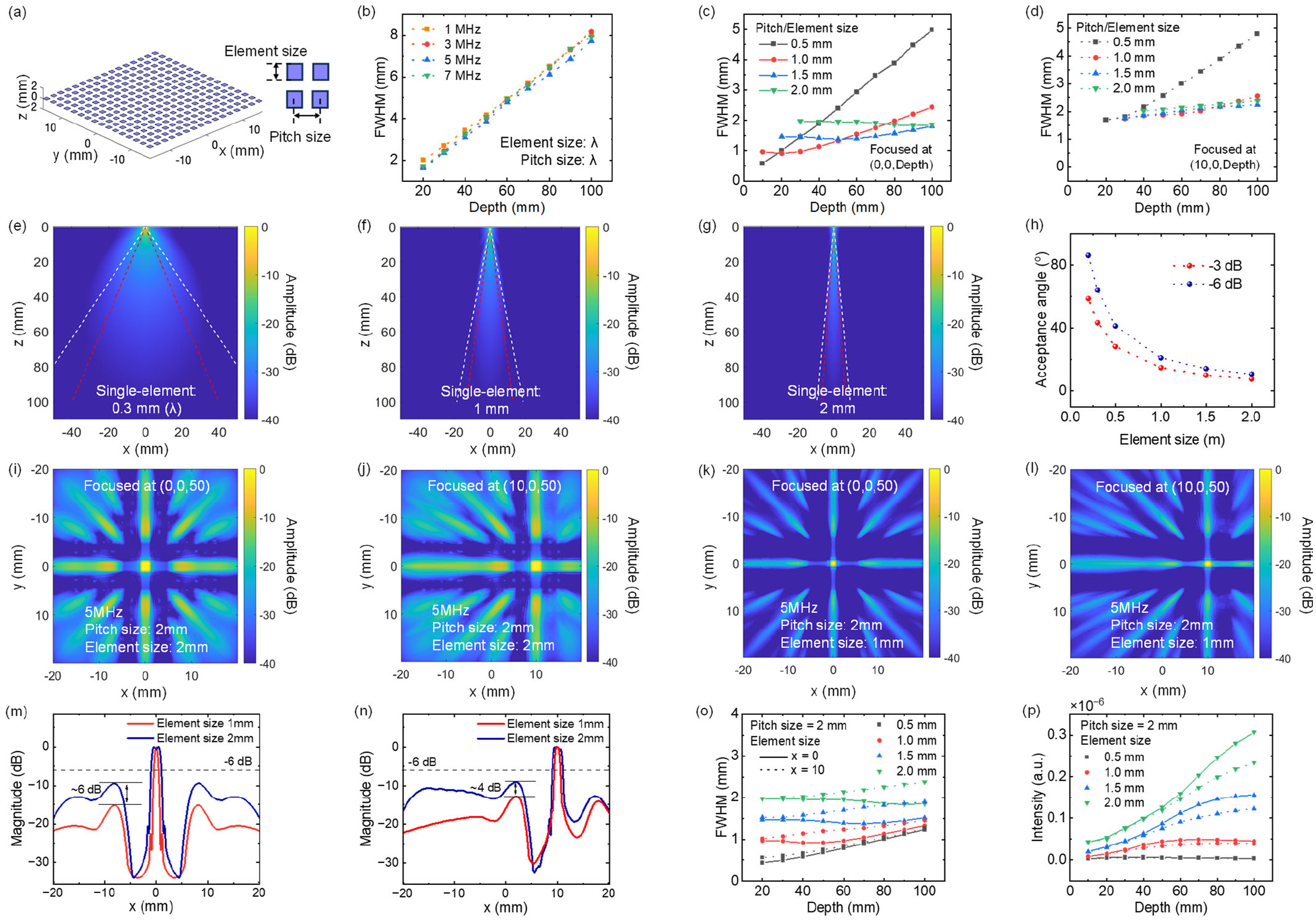
Simulation-guided design of the planar array. (a) Schematic diagram of the array geometry and key design parameters. (b) Full width at half-maximum (FWHM) as a function of depth for conventional planar arrays at different operating frequencies. (c), (d) FWHM as a function of depth for planar arrays with different element and pitch sizes under on-axis focusing and 10-mm off-axis focusing, respectively. (e)–(g) Single-element directivity profiles for element sizes of 0.3 mm, 1 mm and 2 mm, respectively. The white and red dash lines indicate the -6 dB and -3 dB acceptance angles, respectively. (h) Acceptance angle as a function of element size. (i)–(l) 2-D beam profiles in the x-y plane at a depth of 50 mm for a 5-MHz array with a 2-mm pitch and element sizes of 1 and 2 mm, focused at (0,0,50) and (10,0,50), respectively. (m), (n) Corresponding 1-D beam profiles for the 5-MHz array with a 2-mm pitch and element sizes of 1 and 2 mm, respectively. (o)–(p) Depth-dependent FWHM and intensity for the 5-MHz array with a 2-mm pitch and different element sizes under on-axis focusing (solid lines) and 10-mm off-axis focusing (dashed lines).

To address this limitation, we increased the array’s aperture by enlarging the element and pitch sizes simultaneously. Fig. 5c illustrates the effect of pitch and element size on lateral resolution. As these dimensions increase from 0.5 mm to 2 mm, the FWHM becomes more uniform across depth, with a ∼50% improvement at > 50-mm depth compared to the 0.5-mm case. Fig. 5d shows an off-center focusing case, where the focus is steered 10 mm away from the array center. The overall trends were consistent with the on-axis condition, but the slightly increased FWHM indicated an approximately 21% degradation in off-axis lateral resolution.

However, enlarging the element size reduces the acceptance angle of each element, limiting the effective contribution of peripheral array elements, and thereby degrading the lateral resolution. As shown in Figs. 5e–g, the -3 dB and -6 dB acceptance angles decrease from 59° to 8° and from 86° to 11°, respectively, as the element size increases from 0.3 mm to 2.0 mm. To mitigate this trade-off, we further optimized the array by reducing the element size to maintain a broader acceptance angle and thus preserve sufficient resolution across the field of view. Figs. 5i–l show 2D beam profiles in the x-y plane at a 50-mm depth. With the pitch size fixed at 2 mm, reducing the element size from 2 mm to 1 mm results in ∼55% and ∼61% reduction in FWHM with ∼6 dB and ∼4 dB suppression of grating lobes relative to the main lobe for on-axis and off-axis conditions, respectively.

Reducing the element size also lowers transducer sensitivity, as smaller elements exhibit greater impedance mismatch with the 50-Ω front-end electronics[43], [47]. Figs. 5o–p further quantify how the FWHM and peak intensity vary with depth across different element sizes. While smaller elements significantly improve lateral resolution, they result in lower sensitivity. At the same time, the disparity between on-axis and off-axis performance becomes less pronounced. Balancing resolution against sensitivity, we selected a 1-mm element size, which maintained an FWHM of approximately 1 mm across the region of interest while preserving sufficient sensitivity for *in vivo* imaging.

### B. Characterization of the planar ultrasound array

Fig. 6a shows a photograph of the planar array mounted on a robotic arm. Fig. 6b provides an exploded view of the array assembly. After folding the FPCB (Fig. 6c), the array was encapsulated in a 3D-printed housing and secured within a mounting frame along with a water-filled coupling bag. Fig. 6d shows a side view of the array stack, comprising a 1-3 piezoelectric composite topped with high- and low-impedance matching layers. Fig. 6e shows the patterned electrodes on the 1-3 composite and the corresponding element layout. Each element measures 1 × 1 mm^2^ with a 2-mm pitch, yielding a total aperture of 30 × 30 mm^2^. Fig. 6f shows the FPCB routing layout, which enables independent electrical addressing of all 256 elements.

**Fig. 6.**
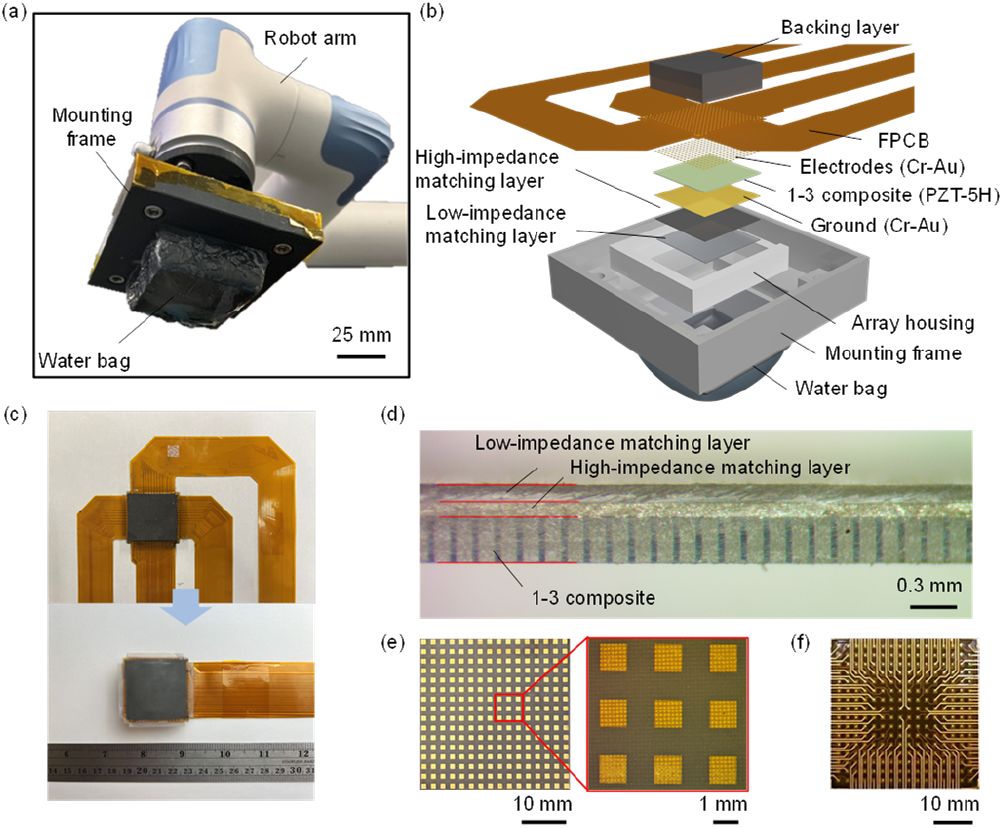
Illustration of planar array for 3D-PCM. (a) Photo of the array mounted on a robot arm. (b) Exploded schematic of the array assembly, including the coupling water bag and mounting frame. (c) Photos of the array before encapsulation. Top: backing-layer side before folding FPCB. Bottom: matching-layer side with folded FPCB. (d) Side-view photo of the dual matching-layer configuration. (e) Photo of the separated 16×16 electrodes for individual element addressing. An enlarged view of the region marked by the red box is shown on the right. (f) Photo of the FPCB bonded to the transducer elements.

Figs. 7a–b show the electrical and acoustic performance of a representative array element. The electrical impedance ranges from 150 to 400 Ω across 4–8 MHz, with multiple resonance peaks arising from the dual matching-layer design, consistent with the KLM simulation. The pulse-echo response shows a center frequency of 5.6 MHz with 72% fractional bandwidth, consistent with the frequency range of the resonance peaks. Figs. 7c, d show the distribution of normalized sensitivity and one-way time-of-flight (TOF) across the array. The mean sensitivity is -2.9 dB with a standard deviation of 2.2 dB, and the TOF root-mean-square error (RMSE) is 0.0149 μs, indicating good element uniformity. Fig. 7e shows the volumetric imaging of the tungsten wire phantom. The close-up views (Figs. 7f–h) demonstrate the array’s capability to resolve two crossing 100-μm tungsten wires located at 125-mm depth. The corresponding spatial resolutions derived from the image are 0.33 mm laterally (x and y directions) and 0.69 mm axially (z direction) (Figs. 7i–j). Fig. 7k presents lateral and axial resolutions as a function of imaging depth. The axial resolution remains ∼0.3 mm beyond 80-mm depth, whereas the lateral resolution degrades from 0.55 to 0.70 mm with increasing depth.

**Fig. 7.**
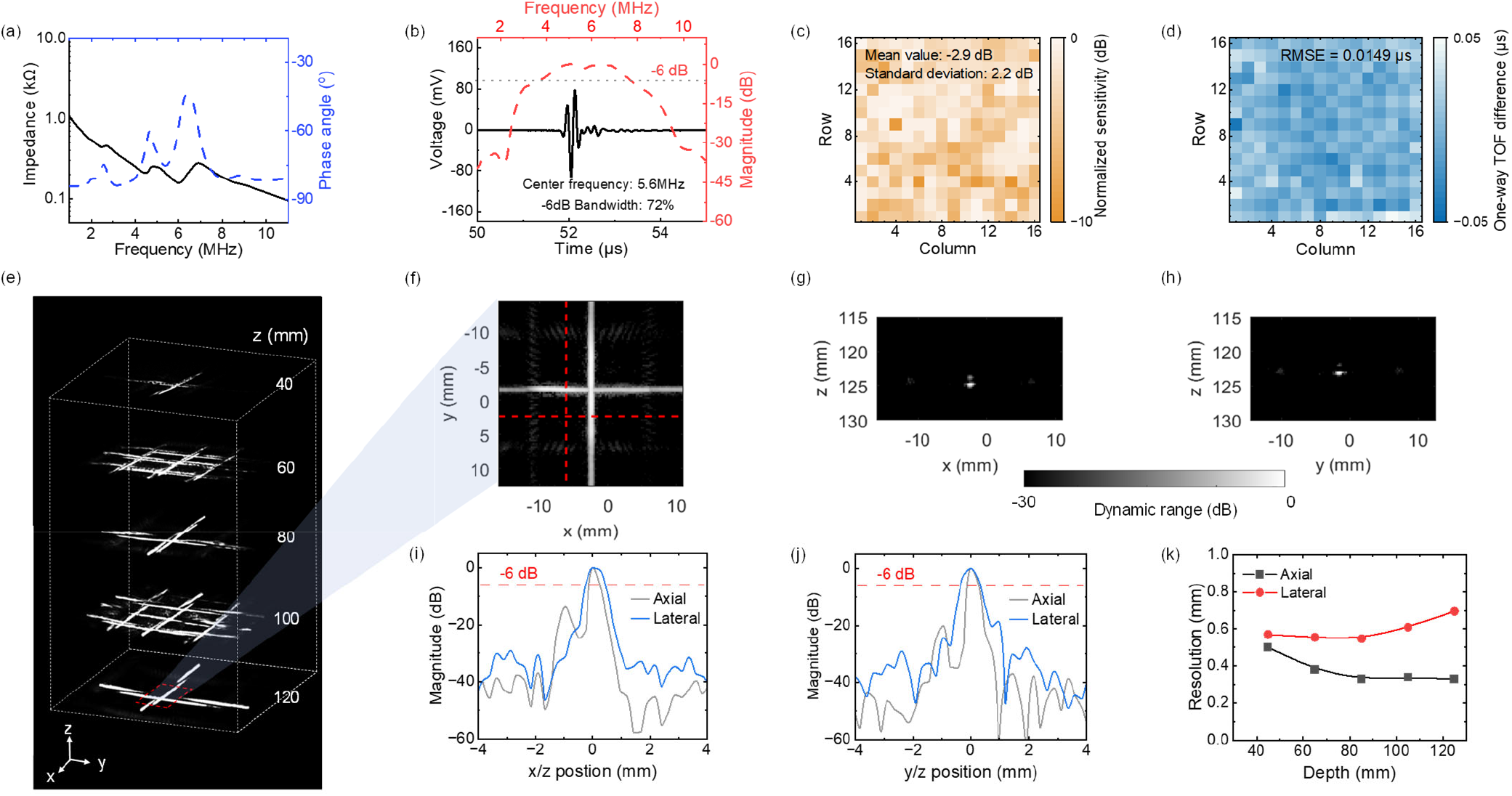
Characterization of the planar array. (a) Electrical impedance magnitude and phase as functions of frequency. (b) Measured pulse-echo response and corresponding frequency spectrum. (c) Normalized sensitivity and (d) one-way time-of-flight (TOF) for individual array element. (e) Volumetric B-mode reconstruction of a 100-µm-diameter tungsten wire phantom. (f) Maximum amplitude projection (MAP) in the x-y plane at the position indicated in (e). (g) x-z and (h) y-z cross-sections at the position marked by the dashed lines in (f). (i) Axial (gray) and (j) lateral (blue) resolution profiles extracted from x-z and y-z cross-sections with -6 dB beamwidths indicated by dashed red lines. (k) Lateral and axial resolutions as functions of imaging depth.

### C. Laser-induced single-bubble cavitation

To validate the performance of the developed planar array, we first examined the cavitation induced by a single laser pulse. Fig. 8a shows the 3D-PCM sensitivity map at various positions relative to the transducer plane, comprising a series of maximum amplitude projection (MAP) images of cavitation events across depths z = 25–105 mm and lateral positions x = 0–20 mm. Distinct bubble clusters were detected at each depth, indicating that our system can image cavitation with high spatial resolution across a large FOV. Representative zoom-in plots of the PCM results at x = 5 mm, y = 0 mm, z = 25 mm are shown in Figs. 8b–e. The top-view map (Fig. 8b) reveals a cavitation cloud with approximately symmetric spread in the x-y plane. Side-view reconstructions along the y-z and x-z planes (Figs. 8c–d) further reveal the confined spatial profile of the cavitation cloud. Fig. 8e shows the volumetric rendering of the cavitation events following a single laser pulse. The model-matrix construction took ∼4 h, and each subsequent PCM image reconstruction took ∼2 min. Fig. 8f shows the PCM signal profiles along the z-axis for five cavitation events at y = 0 mm, x = 5 mm, and z = 25, 45, 65, 86, and 105 mm. Fig. 8g shows the lateral (x-axis) and axial (z-axis) FWHM. These results demonstrate that single-pulse LL-induced cavitation can be imaged in 3D with sub-millimeter resolution using our planar array, providing sufficient spatial precision for the subsequent *in vivo* validation.

**Fig. 8.**
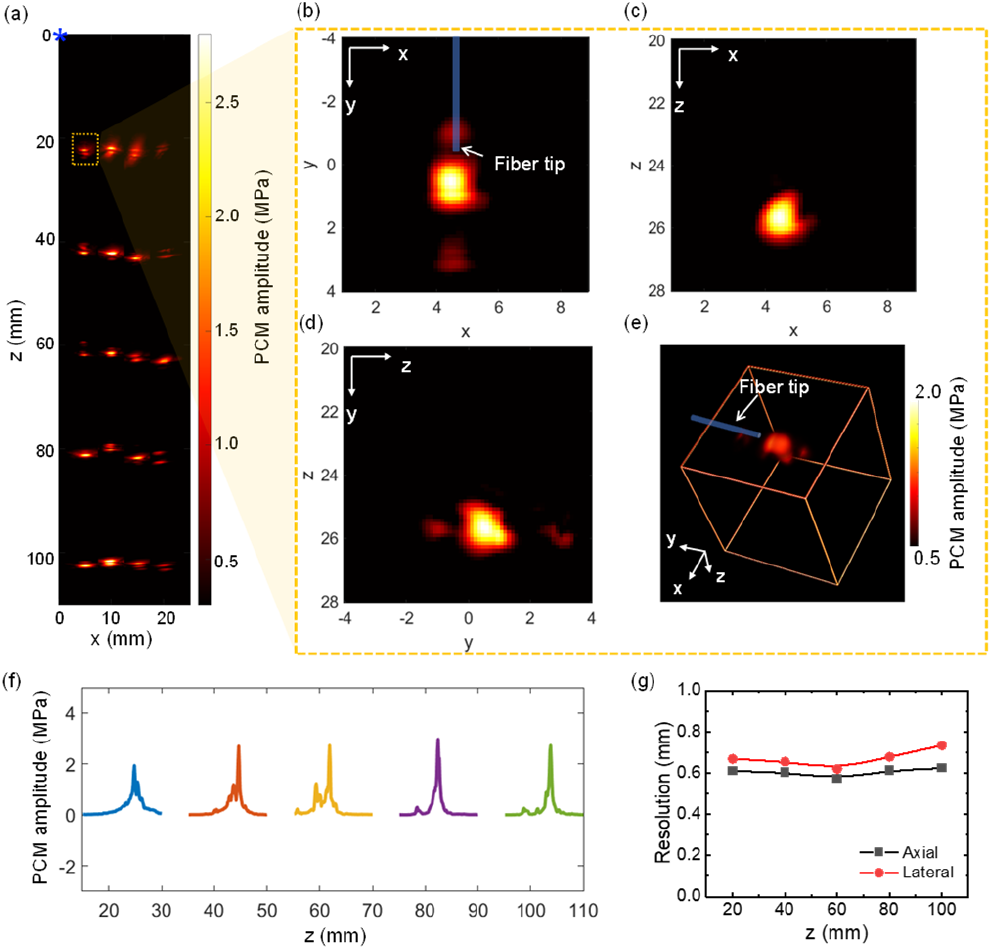
3D-PCM of LL-induced cavitation generated by a single laser pulse. (a) Reconstruction of the PCM sensitivity field (side-view MAP images of the single-pulse-induced cavitation at different positions). The central axis of the transducer array is marked by blue asterisk at (0, 0, 0). The fiber tip was placed parallel to and below the transducer surface at different positions. (b) x-y top-view MAP of the single-pulse induced cavitation at x = 5 mm, z = 25 mm. (c) x-z side-view MAP. (d) y-z side-view MAP. (e) 3D rendering. (f) PCM signal profiles of the single-pulse-induced cavitation at x = 5 mm, y = 0 mm, and z = 25, 45, 65, 85, 105 mm, respectively. (g) Resolution quantification from the single-pulse induced cavitation at x = 5 mm, z = 25, 45, 65, 85, 105 mm.

### D. Validation of 3D-PCM with high-speed camera

To validate 3D-PCM, we compared the 3D-PCM reconstructions with high-speed camera imaging of cavitation dynamics near a glass surface (Fig. 9). The 3D rendering (Fig. 9a) demonstrated an elongated bubble structure extending from the fiber tip, consistent with the anisotropic expansion expected in the presence of a rigid, confined boundary [24], [48]–[50]. Collapse-time analysis (Fig. 9b) showed reproducible intervals of 925–1275 μs across successive events. In the experimental setup (Fig. 9c), the fiber tip was positioned adjacent to the glass surface and imaged with a high-speed camera synchronized with the PCM acquisition. High-speed camera recordings (Fig. 9d) captured the temporal evolution of bubbles following a single laser pulse: an initial cavitation event appeared at 620 μs, followed by a first collapse at ∼950–960 μs and a second collapse at ∼1220–1230 μs. These results confirmed that multiple collapse-rebound cycles occur after a single laser pulse. Correspondingly, the PCM MAP images (Fig. 9e) captured five distinct cavitation events (labeled #1–5), which corresponded precisely to the bubbles observed in the high-speed camera recordings. The spatial localization of each bubble in PCM agreed well with the camera recordings. Together, these findings confirmed 3D-PCM as a reliable approach for volumetric cavitation mapping during LL.

**Fig. 9.**
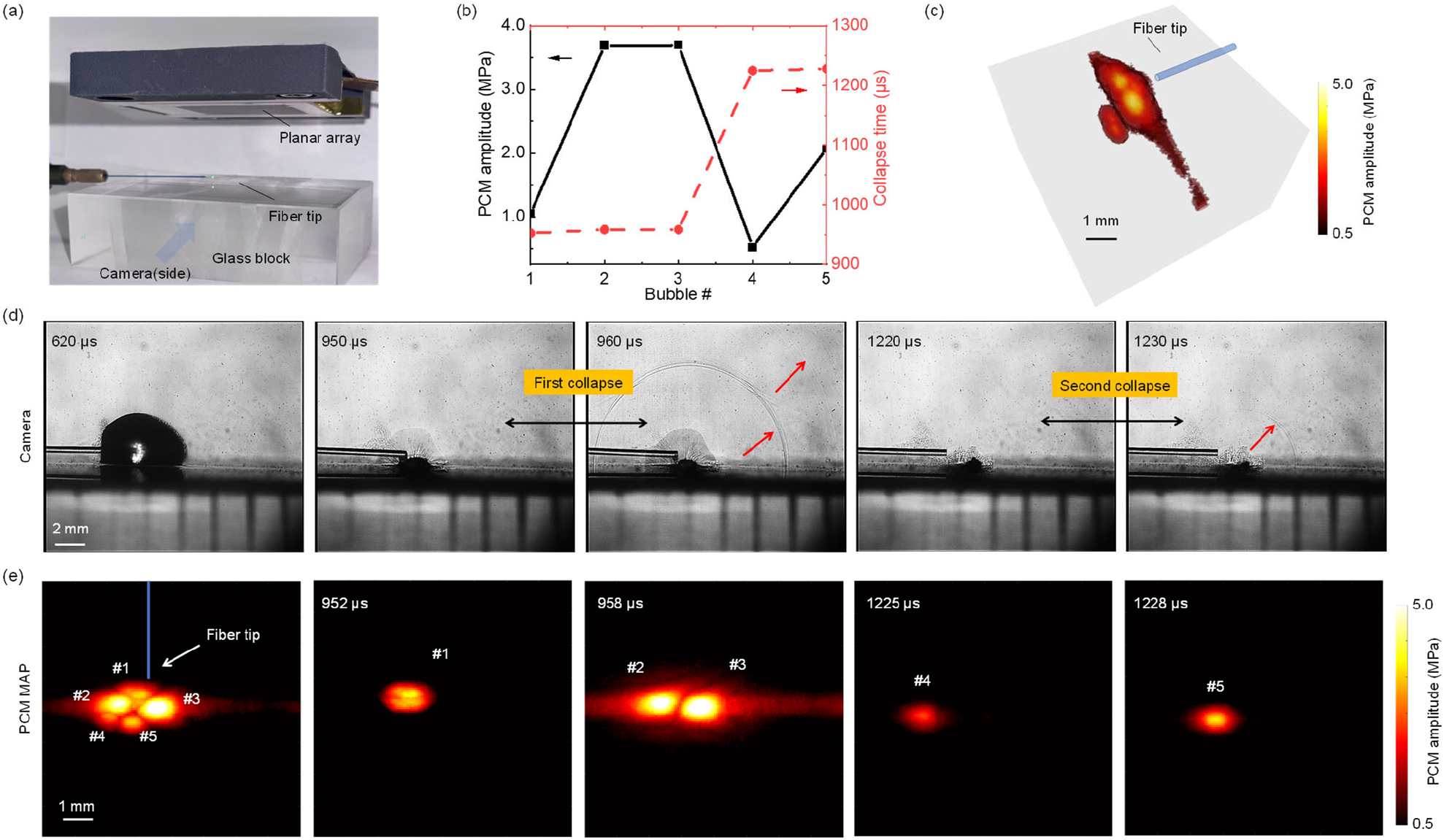
Validation of 3D-PCM using high-speed camera imaging. (a) Photo of experimental setup. (b) Bubble collapse-time analysis. (c) 3D rendering of LL-induced cavitation with a single laser pulse near a glass surface (stand-off distance = 1.0 mm). (d) Side-view high-speed camera images of the first and second bubble collapses. (e) Top-view PCM MAP images of five distinct cavitation events (#1–#5), where the first panel shows the cumulative cavitation MAP, followed by four MAPs showing individual cavitation events in sequence.

### E. In vivo validation of 3D-PCM on a swine model

While benchtop phantom experiments successfully demonstrated the high spatiotemporal resolution of the planar array-based 3D-PCM, the clinical translation of this technology requires validation in a highly scattering and dynamic biological environment. To evaluate the system’s performance under realistic surgical conditions, including anatomical constraints, tissue attenuation, and continuous saline irrigation, we applied the 3D-PCM to monitor cavitation during LL on an *in vivo* swine model, which closely mimics human renal anatomy and clinical workflows (Fig. 10). Endoscopic video recordings (Fig. 10a, top row) confirmed bubble formation near the fiber tip during the treatment. The corresponding PCM images (Fig. 10a, bottom row) show that the cavitation was initially confined around the fiber tip (after ∼600 pulses) and gradually expanded to multiple sites with increasing exposure (1200–3000 pulses). The spatial distribution of cavitation became more heterogeneous at higher pulse counts, reflecting progressive tissue interaction and cumulative cavitation effects.

**Fig. 10.**
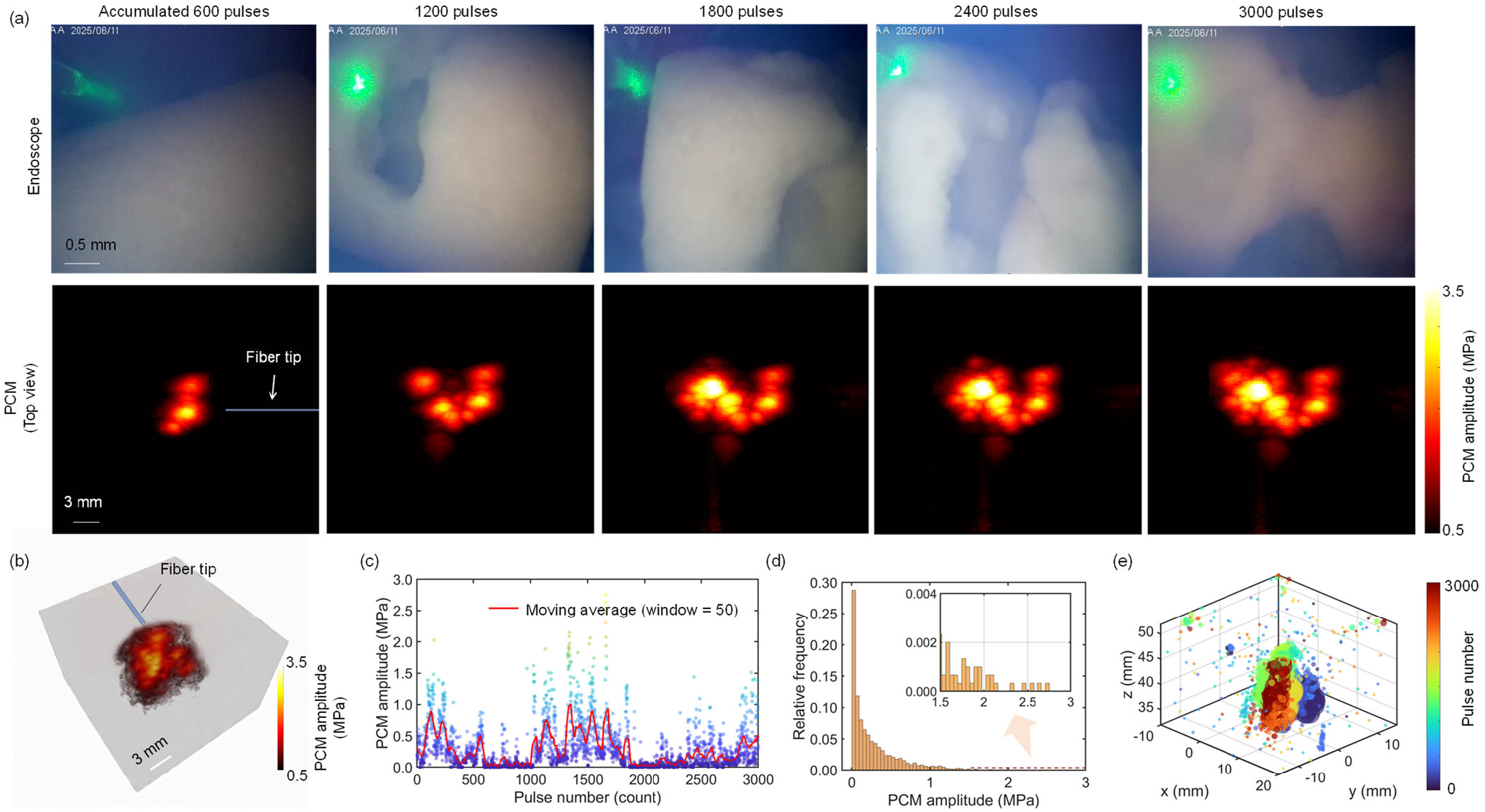
3D-PCM of *in vivo* swine LL treatment. (a) Endoscopic video snapshots (top row) and top-view MAP images of 3D-PCM (bottom row) acquired after 600, 1200, 1800, 2400 and 3000 laser pulses. (b) Volumetric rendering of the accumulated 3D-PCM signal after 3000 laser pulses. (c) PCM peak amplitudes from individual laser pulses with a 50-pulse moving average. (d) Relative-frequency distribution of PCM amplitude. The inset shows the high-amplitude tail. (e) 3-D distribution of cavitation events over 3000 laser pulses, where the scatter size denotes PCM amplitude and the color denotes pulse number.

The volumetric rendering after 3000 pulses (Fig. 10b) revealed an extensive cavitation field extending several millimeters around the fiber tip, with localized high-intensity regions representing repeated bubble collapses. Temporal analysis of PCM signals showed substantial pulse-to-pulse variability in PCM amplitude (Fig. 10c). Although most pulses generated low-amplitude cavitation signals, intermittent high-amplitude events were observed throughout the treatment. This trend was further confirmed by the relative-frequency distribution (Fig. 10d). This finding indicates that LL-induced cavitation is highly dynamic, with transient but intense bubble activity that may contribute to stone fragmentation as well as to collateral tissue effects [46], [47].

The 3D bubble localization analysis (Fig. 10e) further aggregated the estimated collapse positions over 3000 pulses and revealed clustered but spatially dispersed cavitation activity surrounding the fiber tip over several millimeters in all three spatial dimensions. Together, these results demonstrate that LL generates sustained yet highly heterogeneous cavitation in both space and time throughout the treatment. The successful volumetric mapping of these dynamics *in vivo* confirms that the planar array-based 3D-PCM can function effectively alongside standard endoscopic equipment. This demonstrates the system’s practical feasibility for monitoring cavitation in a realistic surgical environment.

## IV. Discussion

We have developed a 3D-PCM system for monitoring cavitation activity during *in vivo* laser lithotripsy. The 3D-PCM system employs a planar ultrasound array with broad bandwidth (∼74%) and element/pitch sizes optimized to balance spatial resolution and sensitivity, enabling detection of bubble collapse events and accurate 3D localization of cavitation within a large field of view (40 × 40 × 110 mm^3^). Compatible with standard clinical workflows, the technology has strong potential for clinical translation as a real-time treatment-guidance tool.

However, the element pitch of our current planar array is much larger than the acoustic wavelength, leading to spatial undersampling and grating-lobe artifacts. Randomizing the element distribution could mitigate these artifacts by spatially averaging the coherent grating-lobe patterns [48]. Alternatively, model-based image reconstruction can suppress the under-sampling artifacts and improve image quality, at the cost of computational time [48], [49]. At present, our reconstruction pipeline operates at ∼1 Hz, which falls short of real-time clinical imaging. Future work will explore algorithmic optimizations to accelerate the reconstruction speed [52]. Furthermore, our planar array can also be extended to photoacoustic tomography (PAT), since its receiving-only imaging process is similar to that of PCM. However, unlike PCM — where the strong ultrasound signals from bubble collapse place low demand on transducer sensitivity — PAT typically requires high receiving sensitivity [49], [50]. For future clinical translation, extensive *in vivo* validation is still needed to quantify the technique’s impact on treatment efficiency. In parallel, we will explore non-imaging-based strategies that leverage raw PCM signals to evaluate treatment effectiveness, thereby relaxing the speed constraints imposed by 3D image reconstruction. This includes non-imaging-based strategies that directly leverage PCM signals to evaluate treatment effectiveness, thereby easing the speed constraints imposed by 3D image reconstruction.

## V. Conclusion

We have developed and validated a 3D-PCM technique based on a customized large-aperture planar ultrasound array, combined with model-based 3D image reconstruction. The system achieves a substantially expanded FOV and imaging depth while maintaining sub-millimeter resolution — all of which are important for clinical translation. Validation with a high-speed camera confirmed the accuracy and robustness of the 3D-PCM system. *In vivo* studies on a swine model showed that the reconstructed cavitation distribution closely correlates with laser-induced stone damage and captures the cumulative, spatially heterogeneous cavitation throughout the procedure [19]. By providing volumetric cavitation feedback with a large FOV during LL, the system has the potential to improve surgical precision, enhance treatment efficiency, and reduce unintended tissue effects [11], [16], [18], [48], [49]. Future work will focus on reducing grating-lobe artifacts by optimizing the array geometry (element pattern) and developing artifact-suppression algorithms, accelerating imaging speed, and exploring signal-based metrics for real-time treatment assessment.

## Acknowledgments

The authors thank the following urologists who were involved in animal experiments and surgeries: David L Barquin, Aaron W Stewart, Nicklas A Sarantos, Jeremy A Kurnot, Logan W Grimaud, Thomas E Schroeder, Megan E Bock, Jiaoti Huang, Jodi Antonelli, Glenn M Preminger, Charles D Scales, Robert A Medairos.

The authors also thank Dornier MedTech for providing the H Solvo laser used in this study. This work was partially sponsored by the United States National Institutes of Health (NIH) grants R01EB037095, RF1 NS115581, R01 NS111039, R01 EB028143, R01 DK139109, R01 DK052985, R01 MH135932; The United States National Science Foundation (NSF) CAREER award 2144788; American Heart Association Collaborative Science Award (25CSA1417550); Duke Gilhuly Acceleration Grant; Duke University Pratt Beyond the Horizon Grant; Eli Lilly Research Award Program; Chan Zuckerberg Initiative Grant (2024-349531); Duke University DST Spark Seed Grant; Duke Coulter Translational Grant; North Carolina Biotechnology Center Triangle Research Grant (2024-TRG-0041).

Dr. C. Qiu gratefully acknowledges Dr. Min Su for providing the matching-layer materials, and Dr. Huaiyu Wu and Dr. Xiaoning Jiang for providing access to the equipment used for transducer characterization.

